# Brain regulatory program predates central nervous system evolution

**DOI:** 10.1101/2021.12.10.472178

**Authors:** Dylan Faltine-Gonzalez, Jamie Havrilak, Michael J Layden

**Affiliations:** Lehigh University, Department of Biological Sciences, Bethlehem, PA USA

## Abstract

Understanding if bilaterian centralized nervous systems (CNS) evolved once or multiple times has been debated for over a century. Recent efforts determined that the nerve chords found in bilaterian CNSs likely evolved independently, but the origin(s) of brains remains debatable. Developing brains are regionalized by stripes of gene expression along the anteroposterior axis. Gene homologs are expressed in the same relative order in disparate species, which has been interpreted as evidence for homology. However, regionalization programs resemble anteroposterior axial patterning programs, which supports an alternative model by which conserved expression in brains arose convergently through the independent co-option of deeply conserved axial patterning programs. To begin resolving these hypotheses, we sought to determine when the neurogenic role for axial programs evolved. Here we show that the nerve net in the cnidarian *Nematostella vectensis* and bilaterian brain are regionalized by the same molecular programs, which indicates nervous system regionalization predates the emergence of bilaterians and CNSs altogether. This argues that shared regionalization mechanisms are insufficient to support the homology of brains and supports the notion that axial programs were able to be co-opted multiple times during evolution of brains.

## Introduction

Efforts to determine if complex brains found in protostome and deuterostome lineages are homologous have been reinvigorated by advances in genomics and phylogenetics. The prevailing model is that brains evolved independently in Xenacoelomorpha ^1–5^, the earliest diverging bilaterian lineage, and that deuterostome and protostome brains are homologous (Figure 1B; Scenario 1). The presence of tripartite brains patterned by strikingly similar mechanisms in *Drosophila* (protostome) and vertebrates (deuterostome) initially argued they arose from a morphologically complex brain in their common ancestor ^6–13^. Fossil evidence and careful mapping of brain morphologies on an improved phylogeny argue that morphological complexity reflects homoplasy rather than homology ^14,15^. While complexity may have evolved multiple times, gene expression patterns along the anteroposterior axis during development of both simple and complex brains appear well conserved (Figure 1A) ^7,16^. Brain-forming regions are regionalized by morphogen gradients with a posterior source of canonical Wnt (cWnt) being ^17–20^. Regionalization generates stripes of gene expression that partition the anteroposterior axis into distinct domains that in turn generate distinct neuronal subtypes (Figure 1A,B). For example, *six3* is a highly conserved anterior region marker that promotes anterior neuronal fates and represses posteriorizing cWnt (Figure 1A) ^21,22^. The remaining regionalized genes (*e*.*g. irx, foxq2, rx, otx, gbx, pax, otp, fez, dlx*) are expressed in similar, albeit not identical, patterns (Figure 1A) ^7–9,11,12,23–36^. Because the regionalization programs in protostome and deuterostome brains are similar, they are thought to have emerged with the brain in their shared ancestor (Figure 1B; Scenario 1). However, investigation of regionalization programs in more species questions whether similar patterning programs truly support homology of protostome-deuterostome brains.

**Figure 1.**
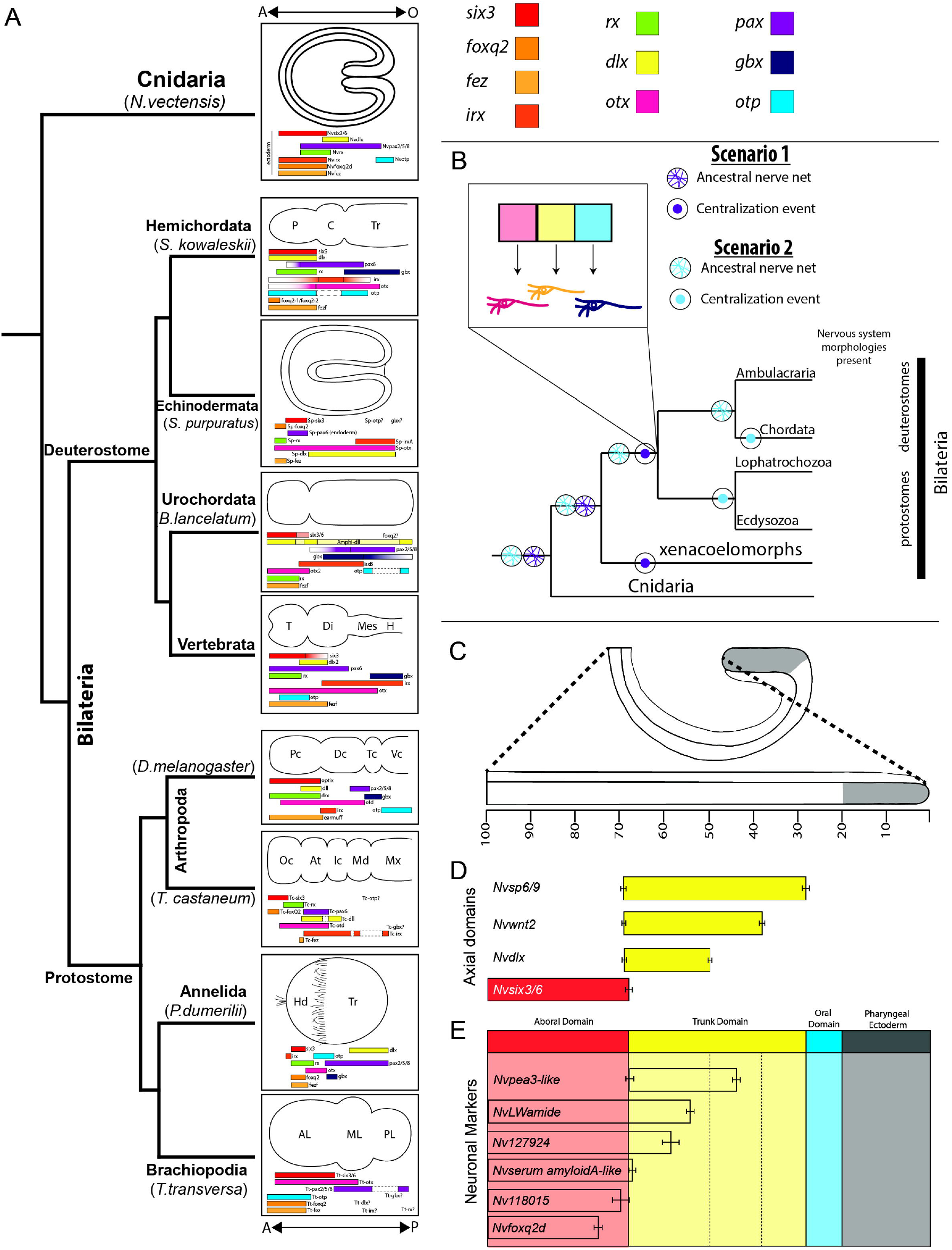
Stripes of regionalized gene expression pattern developing nervous systems. **(A)** Published expression domains of regional gene homologs shown in representative taxa from bilaterian species and *Nematostella*. **(B)** Schematized mechanism by which neural ectoderm is regionalized to pattern neuronal subtypes. **(C)** Positional information was normalized to percent embryo length with 0% being the tip pharyngeal ectoderm (grey) and 100% at the aboral pole. **(D)** Average boundaries of candidate neuronal subtype regulators tested in this study. **(E)** Average expression domain for neuronal subtype markers. The bars indicate the 95% confidence interval for each domain boundary.

Genes that regionalize developing brains function broadly in anteroposterior axial patterning. For example, regionalized genes pattern the entire ectoderm of hemichordates, echinoderms, brachiopods, and cnidarians, all of which lack centralized nervous systems and/or brain-like anterior neuronal condensations (Figure 1A) ^9,12,32,34^. Even within species that have a brain, stripes of regional genes are not restricted to the neuroectoderm^25,35–40^. These data suggest that regionally expressed genes in the developing brain are part of a global anteroposterior axial patterning program rather than a brain specific program. The argument that brain regionalization programs are part of a global axial patterning program, coupled with arguments that the deuterostome ancestor had a nerve net rather than a CNS, has led to the hypothesis that conserved expression of regionalized genes in protostomes and deuterostomes is convergent resulting from co-option of axial programs as brains independently evolved (Figure 1B - Scenario 2). To gain a better understanding of the significance of similar regionalization programs with regards to brain evolution, it is critical to determine when the neurogenic role for axial programs evolved. For conserved regionalization programs to provide strong support for homology, the emergence of their neurogenic role should coincide with the protostome-deuterostome ancestor (Figure 1B).

Cnidarian nerve nets and bilaterian centralized nervous systems evolved from a nerve net-like nervous system in their common ancestor ^41^. Expression data suggests that homologs of bilaterian brain regionalization programs may regionalize the nerve net along the oral-aboral axis of cnidarians. Graded cWnt activity highest at the oral end of the cnidarian sea anemone *Nematostella vectensis* patterns homologs of bilateiran regionally expressed genes into stripes along oral-aboral (O-A) axis using the same regulatory logic observed in bilaterian anteroposterior axis ^42,43^. Moreover, homologs are expressed in the same relative order in both bilaterians and *Nematostella* ^42–47^ (Figure 1A). Although the full complement of *Nematostella* neuronal subtypes is poorly understood, expression of the few known subtype markers indicate that the nerve net is regionalized along the O-A axis during development ^48,49^. Here we test the hypothesis that homologs of the bilaterian anteroposterior regionalization program evolved a role in neuronal patterning prior to the emergence of bilaterians and brains by determining if they pattern neural subtype markers during *Nematostella* development.

## Results

### Identifying regional neuronal subtype specifiers

To identify candidate regulators of neuronal subtypes the expression domain of neuronal markers and regional genes were quantified by calculating the oral and aboral expression limits described as percent embryo length (PEL) (Figure 1C-E; Extended Data 1) (See Materials and Methods). Using this approach we confirmed expression of *Nvfoxq2d* within the *Nvsix3/6+* aboral domain (Figure 1C-E; Extended Data 1A,B) ^50^, and identified *Nv118015* and *Nvserum amyloid A-like* as additional aboral domain neuronal subtypes (Figure 1D-E; Extended Data 1A,C,D). *Nv127924+* and *NvLWamide-like+* neurons are expressed throughout the aboral and the *Nvsp6-9, Nvwnt2*, and *Nvdlx+* trunk domains (Figure 1D,E; Extended Data 1E,F). *Nvpea3-like* initiates at the boundary of trunk and aboral domains and spans the *Nvdlx+* trunk region terminating within the *Nvwnt2+ Nvsp6-9+* domain (Figure 1D,E; Extended Data 1G-J). Because cWnt regulates expression of regionalized genes, the impact of cWnt on neuronal subtype marker expression was tested (Extended Data 2). Reduced cWnt expanded the oral limit of *Nvsix3/6* expression and aboral neuronal subtype expression (Extended Data 2E’,G’,H’)^43^. Similarly, the aboral boundary of *Nvpea3-like* and a subset of trunk regional genes was shifted orally (Extended Data 2A’, D’). Increased cWnt reduced the aboral domain and aboral neuronal subtype expression while expanding the aboral boundary of trunk regional gene and *Nvpea3-like* expression, which was previously observed for trunk markers (Compare Extended Data 2A-H with Extended Data A’’-H’’)^43,46,47^. These data support previous observations that the nerve net is regionalized ^49,50^, identified aboral neuronal subtypes potentially downstream of *Nvsix3/6*, and suggested that *Nvpea3-like* is potentially downstream of trunk identity.

### *Nvsix3/6* specifies aboral neurons

shRNA mediated gene knockdown of *Nvsix3/6* resulted in loss of aboral neurons. Aboral subtype markers *Nvserum amyloid A-like, Nvfoxq2d*, and *Nv118015* were undetectable in many injected animals. When present, their domain was severely reduced (Compare Figure 2A-C with Figure 2A’-C’; Extended Data 3B,B’). *Nv127924* was also reduced in *Nvsix3/6* shRNA injected animals (Extended Data 2A,A’), but *NvLWamide-like* expression was not noticeably impacted by loss of *Nvsix3/6* (Extended Data 3C,C’). Loss of aboral subtype markers was accompanied by an aboral expansion of trunk regional genes and *Nvpea3-like* into the aboral domain, which mimics the *Nvsix3/6* morphant and increased cWnt phenotypes (Figure 2D-F, D’-F’’; Extended Data 2D, D’) ^46,47^. Injection of *Nvsix3/6:venus* mRNA into single cell zygotes resulted in ubiquitous misexpression of *Nvsix3/6* (Figure 2A’’) and expanded the oral expression limit for all neuronal subtypes (Figure B’’,C’’,F’’; Extended Data 3A’’-C’’). Although the oral boundary of *Nvpea3-like* shifted orally, the overall expression of *Nvpea3-like* was severely reduced (Figure 2F’’). *Nvwnt2, Nvdlx*, and *Nvsp6-9* expression was undetectable in *Nvsix3/6* overexpressing animals (Figure 2D’’,E’’; Extended Data 3D’’). These findings suggest that *Nvsix3/6* is necessary and sufficient to promote aboral neuronal fates.

**Figure 2.**
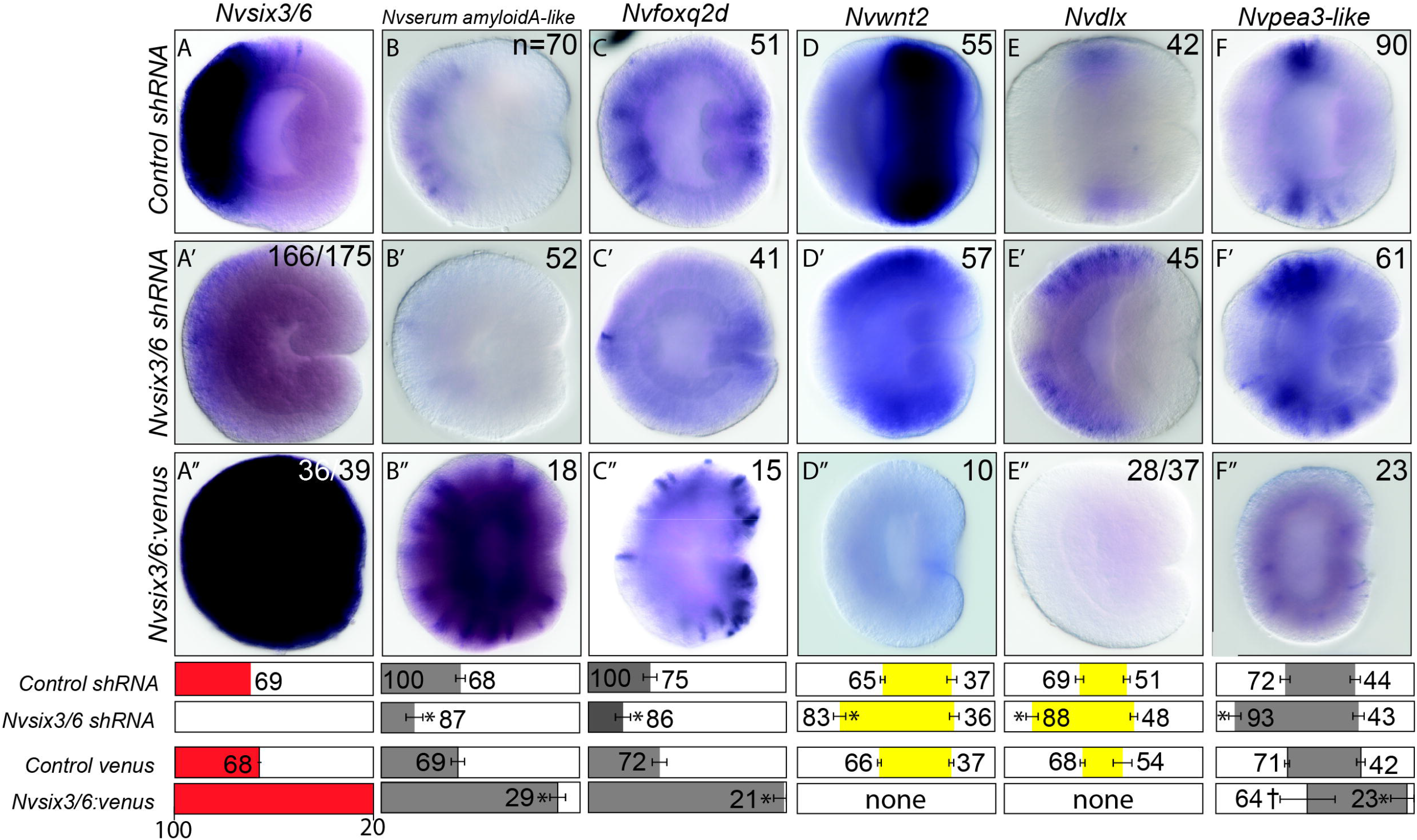
*Nvsix3/6* is *necessary and sufficient to promote aboral fates*. **(A-F)** Control sRNA injected animals. **(A’-F’)** *Nvsix3/6* shRNA injected animals. **(A’’-F’’)** *Nvsix3/6:venus* mRNA injected animals. In all images, aboral is to the left and oral is to the right. Quantification of percent embryo length (PEL) positions for the aboral (left-boundary) and oral (right-boundary) expression limits for each treatment and their control are shown below the images. Bars represent 95% confidence interval. * indicates that treated values are statistically different (p < 0.05) from controls and that the 95% confidence intervals for the values do not overlap, and † indicates that treated values are statistically different (p < 0.05) from controls, but that 95% confidence intervals overlap.

*Nvsix3/6* is known to antagonize cWnt activity in the aboral domain and anterior brain^47^. Thus, it was not clear if *Nvsix3/6*, low levels of cWnt, or both activate aboral neuronal gene expression. To determine if *Nvsix3/6* likely regulates aboral gene expression directly, we screened the previously identified *Nvserum amyloid-A* like and *Nvfoxq2d* enhancers and found three putative *Nvsix3/6* binding sites in each (Figure 3A,C). To test the significance of the *Nvsix3/6* binding sites, wild type (Figure 3A,C) and modified enhancer fragments (Figure 3B,D) driving *mcherry* were injected and the number of positive cells in the aboral domain was quantified in mosaic F0 animals. The wild type fragment had aboral expression in ∼75% of the animals with ∼25% of animals exhibiting medium to high levels of expression (Figure 3A). Deleting the two proximal sites resulted in expression in less than half of the animals, with only 3% of the animals having medium to high expression (Figure 3B). ∼65% of wild type *Nvfoxq2d* enhancers showed expression (Figure 3C), whereas 87% of animals injected with all three *Nvsix3/6* removed had no detectable expression (Figure 3D).

**Figure 3.**
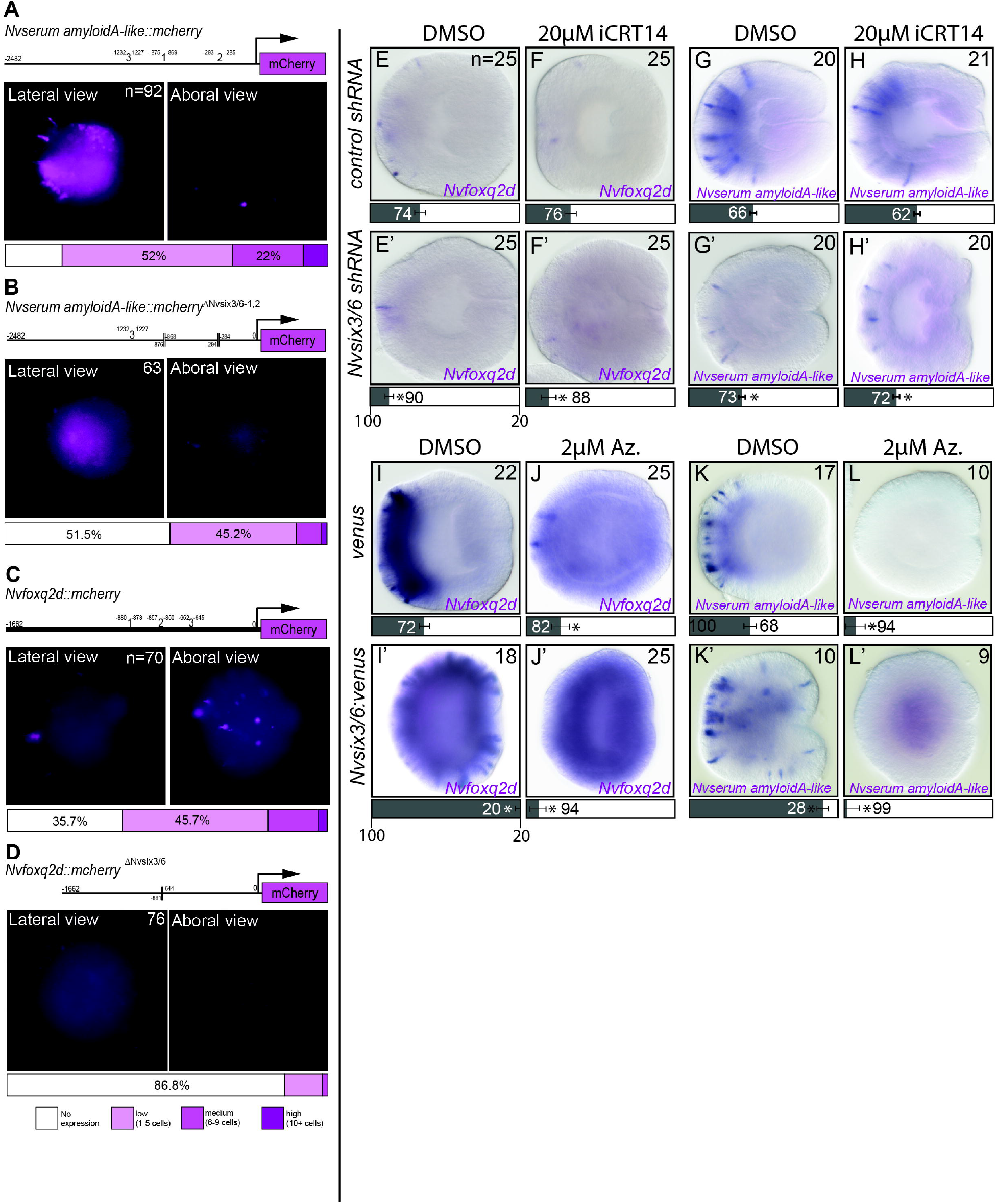
*Nvsix3/6* activates aboral gene expression. **(A-C)** *Nvserum-amyloid A-like::mcherry* transgenes were tested for expression in F0 animals. Wild-type **(A)** shows aboral expression, which is severely reduced in enhancers lacking *Nvsix3/6* binding sites **(B**,**C). (D-G)** The impact of iCRT14 treatments on aboral gene expression in uninjected animals **(E**,**G)** or in animals injected with *Nvsix3/6* shRNA **(E’**,**G’)** compared to controls **(D-F). (H-K)** The impact of Azankenpaullone treatments on aboral gene expression in uninjected animals **(I**,**K)** or in animals injected with *Nvsix3/6* mRNA **(I’**,**K’)** compared to controls **(H-J)**. Bars represent 95% confidence interval. * indicates that treated values are statistically different (p < 0.05) from controls and that the 95% confidence intervals for the values do not overlap, and † indicates that treated values are statistically different (p < 0.05) from controls, but that 95% confidence intervals overlap.

We reasoned that if low cWnt levels played a significant role in promoting aboral neuronal fates, then treating *Nvsix3/6* knockdown animals with the cWnt antagonist iCRT14 should rescue the reduction of aboral neuronal fates. Treatment of *Nvsix3/6* shRNA injected animals with 20µM iCRT14 did not rescue either *Nvserum amyloid A-like* or *Nvfoxq2d* expression (Figure 3F,F’-H,H’). To determine if suppression of cWnt is required to specify aboral fates, *Nvsix3/6* misexpressing animals were treated with 2µM of the Wnt agonist Azankenpaullone. Misexpression of *Nvsix3/6:venus* failed to rescue aboral fates lost by treatment with Azankenpaullone (Figure 3J,J’,L,L’). Loss of aboral subtypes was accompanied by an expansion of the *Nvpea3-like* into the aboral domain, which was also not rescued by *Nvsix3/6* misexpression (Extended Data 4B,B’). Overall, the phenotypes resembled the Azankenpaullone only treatments (Compare Figure 3J’ and L’ with Extended Data 2G” and H”) suggesting that either high levels of cWnt, a trunk regional marker activated by cWnt, or both repress aboral neuronal fates. Collectively these findings suggest that *Nvsix3/6* promotes aboral fates by activating expression of aboral neuronal subtype markers and by inhibiting trunk identity through repression of cWnt.

### *Nvwnt2* promotes trunk and represses aboral neuronal subtype identity

To gain insights about the neurogenic role of trunk genes, we reduced *Nvwnt2* and *Nvdlx* with shRNA mediated knockdown (Figure 4). Loss of *Nvdlx* did not disrupt expression of regional genes or neuronal markers along the O-A axis (Extended Data 5A,A’-H,H’). Reduction of *Nvwnt2* resulted in an oral shift in the aboral boundary of *Nvsp6/9* and *Nvdlx* (Figure 4A,A’-C,C’). The aboral boundary of *Nvpea3-like* expression was also shifted orally, and the overall level of *Nvpea3-like* expression was reduced (Figure 4D,D’). The reduced levels of *Nvpea3-like* resemble the reduced levels observed in iCRT14 treated and *Nvsix3/6* overexpressing embryos (Extended Data 2D’; Extended Data 4A’, respectively). Although it should be noted that it is not clear if *Nvwnt2* is capable of activating the cWnt pathway^51^. Loss of *Nvwnt2* resulted in an oral expansion of the aboral domain gene *Nvsix3/6* (Figure 4E,E’) and the neuronal subtype markers *Nvfoxq2d* and *NvLWamide-like*, but not *Nvserum-amyloid A-like* (Figure 4F,F’-4H,H’). *NvsoxB(2)* and *Nvath-like*, neural progenitor markers^52,53^, were not reduced in *Nvwnt2* shRNA injected animals (Extended Data 5I) indicating that loss of *Nvpea3-like* is not due to a reduction in neuronal progenitors but rather from changes to patterning. These observations suggest *Nvwnt2* promotes the trunk neuronal gene *Nvpea3-like* and represses the oral boundary of *NLWamide-like* in the trunk and the aboral *Nvfoxq2d* neurons.

**Figure 4.**
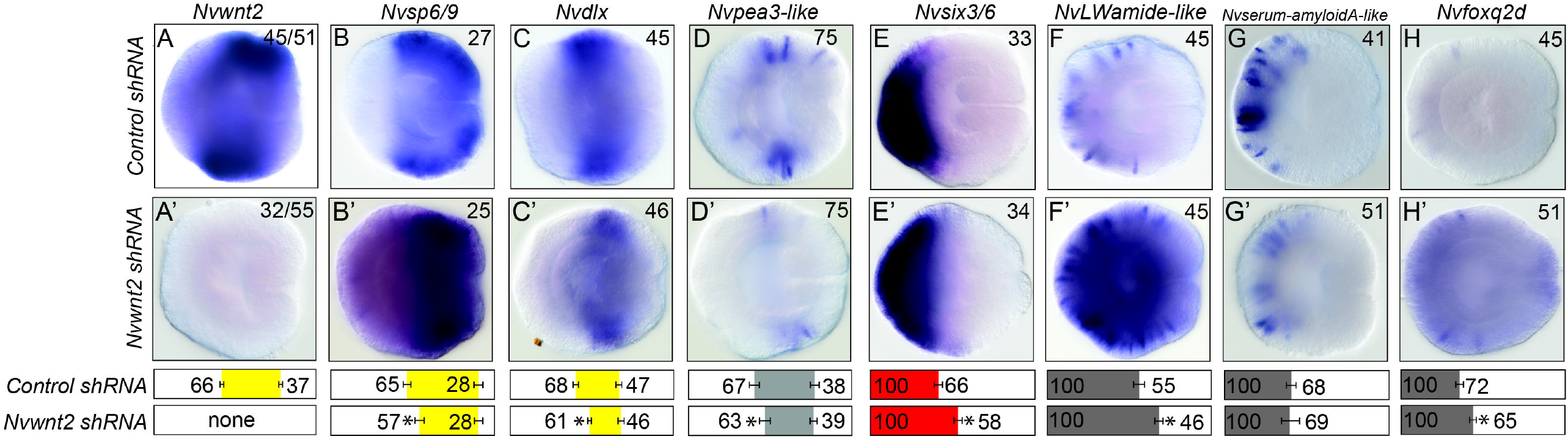
*Nvwnt2* promotes trunk identity and suppresses aboral fates. **(A-H)** Control sRNA injected animals. **(A’-H’)** *Nvwnt2* shRNA injected animals. In all images, aboral is to the left and oral is to the right. Quantification of percent embryo length (PEL) positions for the aboral (left-boundary) and oral (right-boundary) expression limits for each treatment and their control are shown below the images. Bars represent 95% confidence interval. * indicates that treated values are statistically different (p < 0.05) from controls and that the 95% confidence intervals for the values do not overlap, and † indicates that treated values are statistically different (p < 0.05) from controls, but that 95% confidence intervals overlap.

### A model for nerve net regionalization along the O-A axis

We propose that during development, neuronal fates are patterned along the O-A axis by stripes of regionalized gene expression that are established in part by graded cWnt activity at the oral end, which is opposed by *Nvsix3/6* and FGF activity at the aboral end ^42,43,46,47^. Regionally restricted domain genes both promote neuronal fates born within their domain, and prevent expression of adjacent domain genes and neuronal fates (Figure 5A). Specifically, we show that the aboral domain gene *Nvsix3/6* promotes aboral fates by activating expression of *Nvserum amyloid A-like* and *Nvfoxq2d* while simultaneously repressing trunk identity. Similarly, within the Trunk region, *Nvwnt2* promotes the expression of *Nvpea3-like* and represses *Nvsix3/6* and *Nvfoxq2d* aboral fates. Additional factors must regulate trunk patterning, as there are a number of regionally expressed genes throughout the trunk that likely generate multiple molecularly distinct domains^43^. Thus, efforts to identify neuronal fates downstream of regionally expressed genes will be necessary to fully describe nerve net patterning. In addition to inputs from regionally expressed genes, the level of cWnt may also influence neuronal patterning directly. For example, *Nvpea3-like* and *NvLWamide-like* are disrupted by pharmacological manipulation of cWnt activity and in animals with reduced *Nvwnt2*. However, we cannot rule out that cWnt is acting through a yet identified regional gene(s).

**Figure 5.**
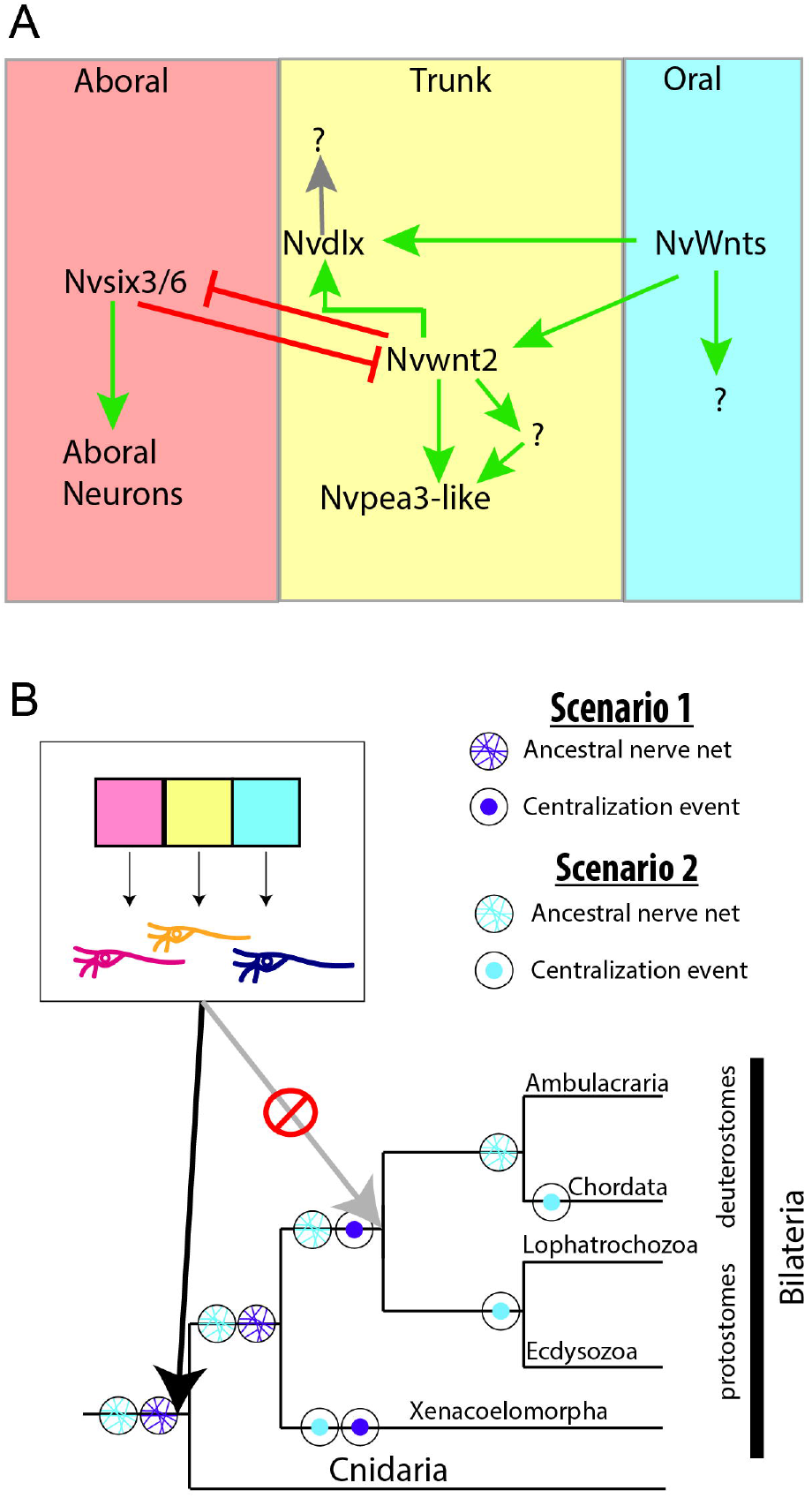
Model of *Nematostella* nerve net patterning and evolution of neurogenic regionalization programs. **(A)** Schematic describing how regionalized genes pattern the developing *Nematostella* nerve net. **(B)** Regional patterning of nervous systems evolved in the cnidarian-bilaterian ancestor, not within the bilaterians.

## Discussion

The mechanisms that regionalize bilaterian brains predate bilaterian divergence and brain evolution. The conserved order for stripes of gene expression observed in bilaterians is also observed in *Nematostella*. In both lineages, regionalized genes regulate neuronal fates born within their respective domains. Some bilaterian regionally expressed genes maintain stripe boundaries through cross-repressive interactions between adjacent genes, which we also observed in *Nematostella*. The similarities in *Nematostella* nerve net and bilaterian brain patterning along the O-A and A-P axes argue that they stem from a shared ancestral program (Figure 5B). Whether brains evolved once or multiple times, they emerged from a regionalized ancestral nerve net. Because regionalization programs broadly pattern the ectoderm in cnidarians and a number of bilaterians, it is reasonable to expect that the stripes of regionalized gene expression would be maintained in the shift to centralization from a nerve net. Especially considering that the tissue that gives rise to brains/CNSs is induced as part of the ectoderm that is later internalized to generate the brain and nerve cords. It is then expected that regionalized gene expression would also be similar whether co-opted once or multiple times as long as the brain is patterned along the anteroposterior axis. These findings reject the argument that similar regionalization programs are sufficient to support the homology of protostome and deuterostome brains.

The pre-bilaterian origin for nervous system regionalization increases support for the co-option hypothesis as no novel function would need to evolve for axial programs co-opted into independently evolved brains. Evidence that co-option occurred independently in deuterostomes and protostomes would be strengthened by confirming that the vertebrate CNS evolved independently. Deuterostomes are comprised of chordates and ambulacrarians (Figure 1A,B). Ambulacrarians lack brains and CNSs and have a nerve net-like architecture ^54^. The mobile predatory lifestyle of ambulacrarian species is inconsistent with secondary loss of centralization. This coupled with the shared basiepidermal plexus found in vertebrates suggested that the ancestral state of deuterostomes was a nerve net resembling the likely architecture of the urbilaterian, which would suggest centralization in chordates evolved independently. Deuterostome regionalized gene expression is regulated by three organizing centers, the ANR, ZLI, and IsO that are deuterostome specific and pattern regionalized genes in the developing vertebrate brain and the anterior ectoderm of the ambulacrarian hemichordate *S. kowalevskii* ^55^. Except the ANR the organizers are deuterostome specific, indicating that unique regulatory logic upstream of regionalized genes evolved coincidently with the stem deuterostome, likely as part of the broad axial patterning program. If a nerve net is the ancestral in deuterostomes, the vertebrate brain/CNS independently evolved and must have co-opted the deuterostome axial programs. As the bilaterian phylogeny is resolved and the neural architecture is described in more species, some hypotheses are emerging that centralization of the nervous system may have also occurred multiple times in protostomes^8,56–59^. Determining if there are unique molecular mechanisms that pattern regional genes at nodes that may have possessed a nerve net, and in the brains of species that stemmed from those nodes may provide insights about the number of times co-option may have occurred. Similarly, because centralization/condensation of the nerve net into nerve cords and a brain occurred independently in the Xenacoelomorpha, efforts should be made to carefully map the regulatory programs within that clade. If regionalized genes patterned the ancestral xenacoelomorph nerve net and later evolving CNSs, it would provide a definitive example of co-option of axial programs into Brain and/or CNS patterning, which would provide compelling support for the co-option hypothesis.

## Acknowledgements

We would like to thank Layla Al-Shaer for comments and suggestions. We also thank Brooke Schaeffer, Caroline Jennings, and Krishna Patel, for their contributions. This research was supported by grants from the National Science Foundation (NSF CAREER 1942777) and NIH National Institute for General Medical Sciences (R01GM127615)

## Figure Legends

**Extended Data 1. Example expression patterns of genes used in this study. (A-F)** Aboral genes. **(G-J)** Trunk genes.

**Extended Data 2. Manipulations of cWnt levels disrupt regional domain gene and neuronal subtype expression patterns. (A-H)** DMSO treated control. **(A’-H’)**

Treatment with 20µM iCRT14 from late blastula to gastrula stage. **(A’’-H’’)** Treatment with 2µM Azenkenpaullone from late blastula to gastrula stage. In all images, aboral is to the left and oral is to the right Quantification of percent embryo length (PEL) positions for the aboral (left-boundary) and oral (right-boundary) expression limits for each treatment and their control are shown below. Bars represent 95% confidence interval. * indicates that treated values are statistically different (p < 0.05) from controls and that the 95% confidence intervals for the values do not overlap, and † indicates that treated values are statistically different (p < 0.05) from controls, but that 95% confidence intervals overlap.

**Extended Data 3. Phenotypes resulting from disruption of *Nvsix3/6*. (A-D)** Control sRNA injected animals. **(A’-D’)** *Nvsix3/6* shRNA injected animals. **(A’’-D’’)** *Nvsix3/6:venus* mRNA injected animals. In all images, aboral is to the left and oral is to the right. Quantification of percent embryo length (PEL) positions for the aboral (left-boundary) and oral (right-boundary) expression limits for each treatment and their control are shown below images. Bars represent 95% confidence interval. * indicates that treated values are statistically different (p < 0.05) from controls and that the 95% confidence intervals for the values do not overlap, and † indicates that treated values are statistically different (p < 0.05) from controls, but that 95% confidence intervals overlap.

**Extended data 4. *Nvpea3-like* expression in Azenkenpaullone treated and *Nvsix3/6* injected animals**. Injection of *Nvsix3/6* mRNA is insufficient to rescue aboral expansion of *Nvpea3-like* in Azankenpaullone treated animals. Bars represent 95% confidence interval. * indicates that treated values are statistically different (p < 0.05) from controls and that the 95% confidence intervals for the values do not overlap, and † indicates that treated values are statistically different (p < 0.05) from controls, but that 95% confidence intervals overlap.

**Extended data 5. *Nvdlx* does not significantly disrupt axial patterning or neuronal subtype patterning**. Control sRNA injected animals. **(A’-H’)** *Nvdlx* shRNA injected animals. **(I)** qPCR in *Nvwnt2* shRNA injected animals. In all images, aboral is to the left and oral is to the right. Quantification of percent embryo length (PEL) positions for the aboral (left-boundary) and oral (right-boundary) expression limits for each treatment and their control are shown below images. Bars represent 95% confidence interval. * indicates that treated values are statistically different (p < 0.05) from controls and that the 95% confidence intervals for the values do not overlap, and † indicates that treated values are statistically different (p < 0.05) from controls, but that 95% confidence intervals overlap.

**Extended Data 6. Table of primers used to generate shRNAs and enhancer constructs used in this work**.

**Extended Data 7. Tables of all raw data quantified in this study**.

## Materials and Methods

### Animal Care, microinjection, and fixation

*Nematostella* polyps were maintained in *Nematostella* media (12ppt artificial sea water (Instant Ocean)), maintained in the dark at 17°C, and fed artemia five times per week. One week prior to spawning induction polyps were fed oyster and the *Nematostella* media was replaced. Spawning was induced as previously described^60^. Microinjections were performed as previously described on a Nikon SMZ1270 stereo scope^61^. Embryos were raised at either 17°C or 22°C to the desired stage. Embryos were fixed and stored as previously described ^62^.

### shRNA, mRNA, pharmacological treatments, and plasmid injections

shRNAs were designed and synthesized as previously described ^63^. shRNA sequences and primers used to generate them can be found in (Extended Data 6) shRNAs were then stored at −80 in single use aliquots. A previously published scrambled sequence shRNA was used as the control for all shRNA injections ^63^. Gene knockdown was confirmed through *in situ* hybridization and/or qPCR. *Nvsix3/6:venus* was generated by subcloning the *Nvsix3/6* coding sequence into pENTR/D TOPO (ThermoFisher Scientific) using published primers previously used to PCR amplify *Nvsix3/6* and synthesize *Nvsix3/6* mRNA ^47^. A 3’ Venus tag was added by recombining the *Nvsix3/6* coding sequence into pSPE3-R-Venus destination vector using the Gateway LR cloning reaction (ThermoFisher Scientific). mRNA synthesized and injected at 300ng/µL using previously described methods ^61^. Pharmacological treatments were performed either from 3 hours post fertilization (hpf) until 24hpf at 22°C or from 24hpf to 48hpf at 17°C. Stocks of the Wnt agonist 1-Azakenepaullone (Sigma A3734) or Wnt antagonist iCRT14 (Sigma SML0203) were generated by dissolving each compound in DMSO at 10mg/mL. Control embryos were treated with DMSO equal to the volume of stock compounds added to the *Nematostella*medium. Embryos were washed with fresh *Nematostella* media prior to fixation.

To synthesize of the *Nvserum amyloid A-likeΔNvsix3/6::mcherry* and the *Nvfoxq2dΔNvsix3/6::mcherry* the predicted Nvsix3/6 binding motifs were first identified using http://cisbp.ccbr.utoronto.ca/TFTools.php and the published binding domain from Sebés et al. 2018 in the known enhancer sequences of *Nvserum amyloid A-like* and *Nvfoxq2d* ^48,50,64^. To remove the predicted *Nvsix3/6* binding sites 1 and 2 in the *Nvserum amyloid A* promoter, the enhancer sequences upstream and downstream of predicted binding sites were subcloned into pGEM-T and the enhancer was reconstituted with binding sites absent (Extended Data 6). To remove the predicted *Nvsix3/6* binding sites in the *Nvfoxq2d* primers were designed to remove a 279bp region containing all three *Nvsix3/6* binding domains (Extended Data 6). Each enhancer, *Nvserum amyloid A-likeΔNvsix3/6* and *Nvfoxq2dΔNvsix3/6*, was then subcloned into the pNvT-mcherry reporter construct using the PacI and AscI restriction sites to place the enhancer upstream of the *mcherry* coding sequence. Plasmids were then injected into wildtype embryos at 60ng/µL. F0s were lightly fixed, as previously described^65^, quantified by recording the number of mCherry positive cells at the aboral end at the late gastrula stages.

### In situ hybridization, imaging, and domain quantifications

*In situ* hybridization was performed using previously published methods^62^. DIC images of *Nematostella* embryos were taken on a Nikon NTi with a Nikon DS-Ri2 color camera using the Nikon Elements software. To quantify domain size, embryos were rotated so that lateral images were acquired at a medial focal plane that allowed identification of pharyngeal ectoderm. Images were then uploaded to Fiji where domain size was measured using the segmented line tool ^48,50,64,66^. Three measurements were taken to determine domain size: (1) total size of embryo from pharyngeal ectoderm to future apical tuft (2) pharyngeal ectoderm to oral most gene expression (3) future apical tuft to the most aboral end of gene expression. We then used these three measurements to determine the start and end of the expression domains along the oral-aboral axis of the embryo, recorded as a percentage of the embryo. The start of the expression domain was calculated by dividing measurement 2 by measurement 1, which was then multiplied by 100. The aboral end of the expression domain was calculated by dividing measurement 3 with measurement 1, multiplying the value by 100, and then subtracted by 100 which demarcates the end of the expression domain.

### Statistical Analysis

Statistical analyses were performed with Microsoft excel (version 16). Data are presented here as the mean value calculated with error bars representing a 95% confidence interval. All raw data are available in (Extended Data 7). Statistical significance was calculated using a two-tailed student t-test assuming unequal variance. The reported n represents the total number of embryos assessed, but experiments included were repeated a minimum of 2 times with all replicates showing the same results. All raw data available in Extended Data 7

## Notes

### Competing Interest Statement

The authors have declared no competing interest.

