## Supplementary figures and images for "Brain regulatory program predates central nervous system evolution"

### Extended Data 1

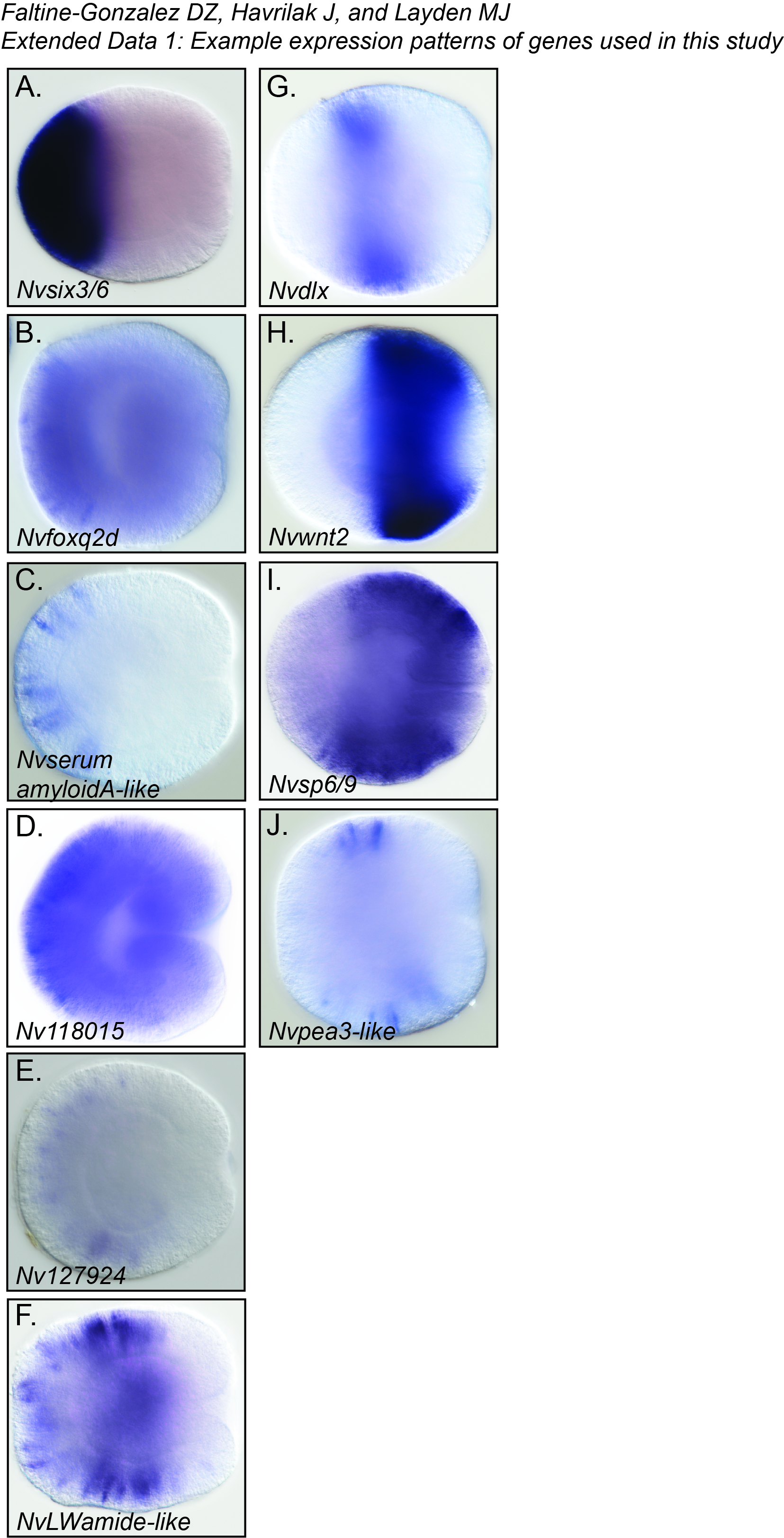

### Extended Data 2

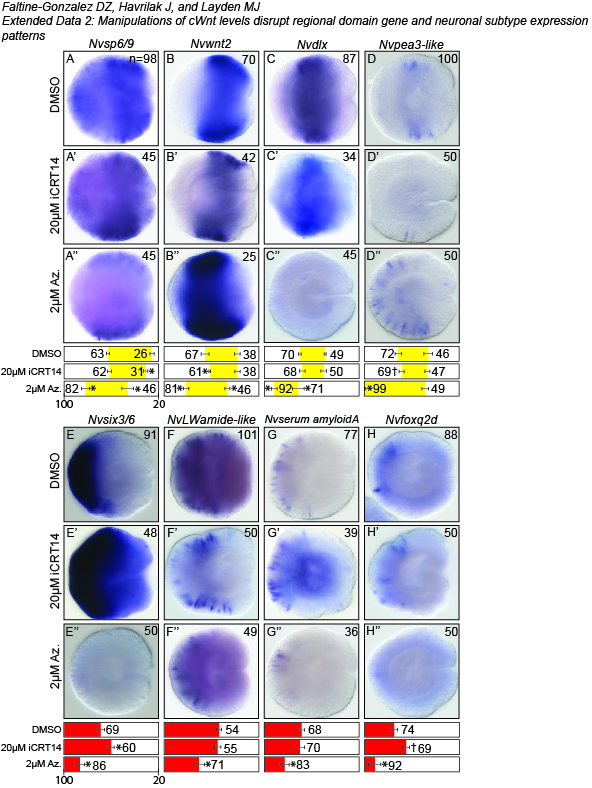

### Extended Data 3

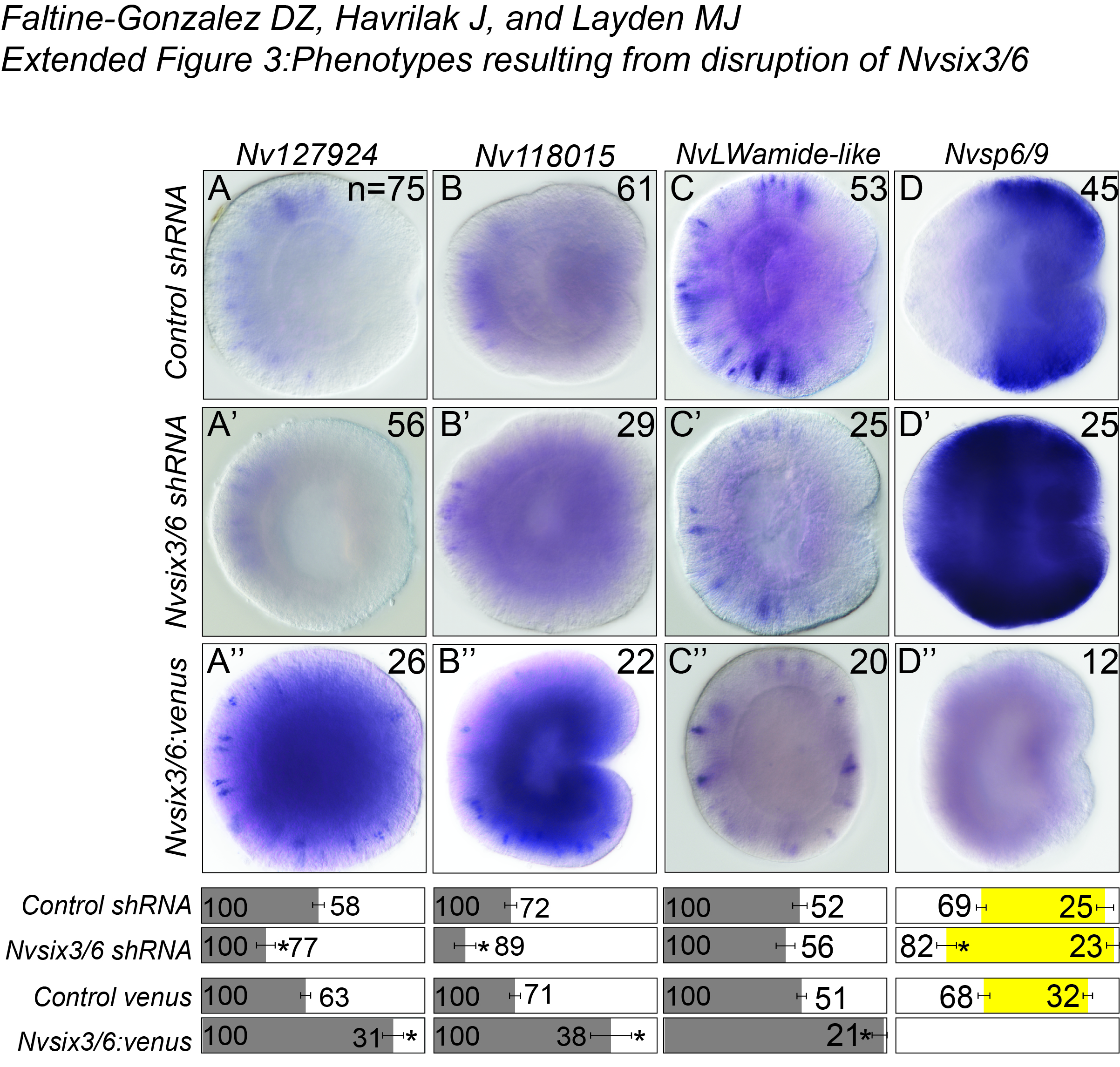

### Extended Data 4

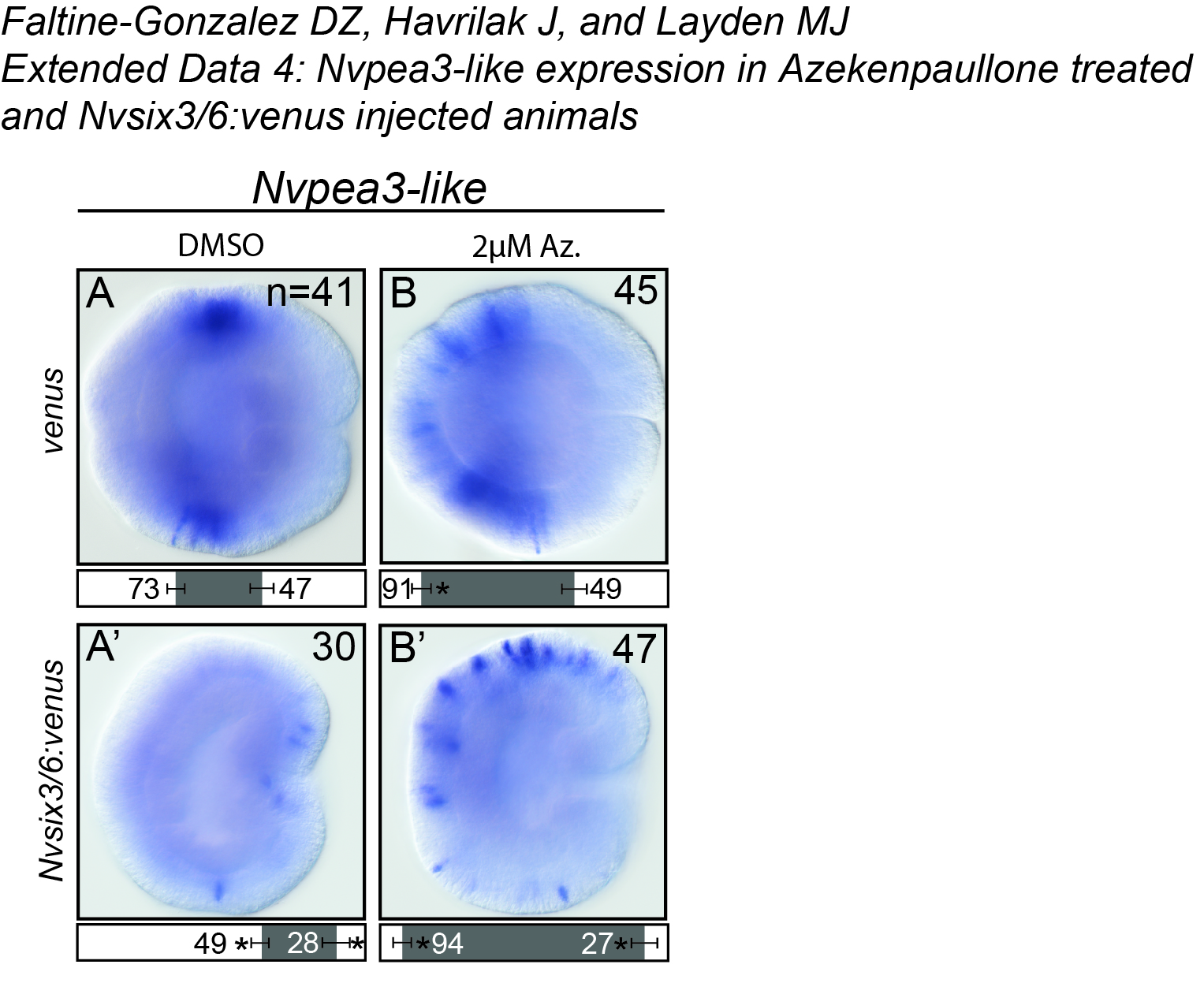

### Extended Data 5

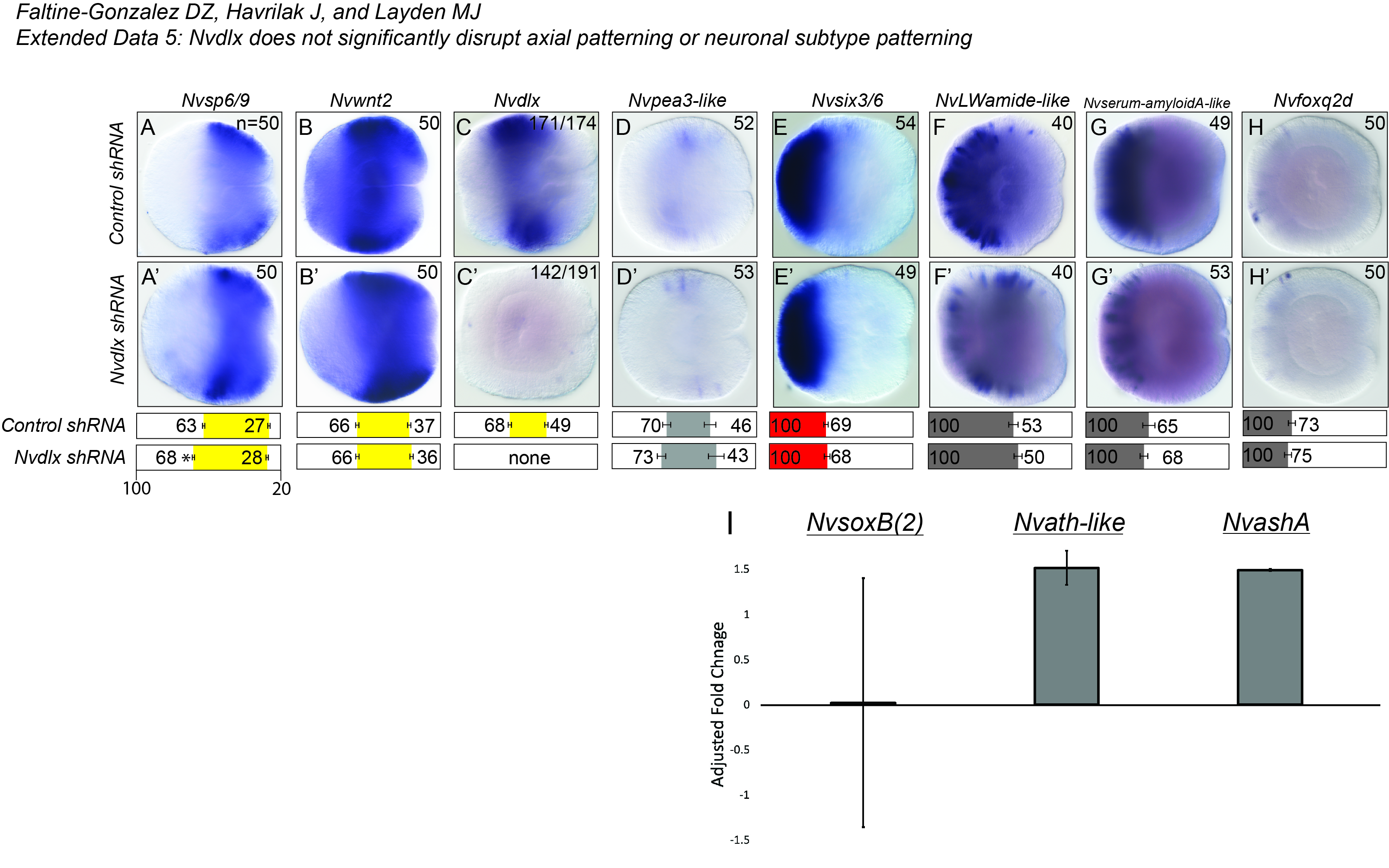
